# Production of unreduced microspores in Arabidopsis flowers cultivated in culture medium suggests a role of sucrose in facilitating meiotic cytokinesis

**DOI:** 10.1101/2025.07.31.667867

**Authors:** Huiqi Fu, Yuting Chen, Xueying Cui, Huishan He, Jingru Wang, Chong Wang, Ziming Ren, Bing Liu

## Abstract

Live-imaging microscopy technology has been increasingly applied for meiosis study in plants, which largely relies on the set up of a healthy *ex vivo* culture system for inflorescences ensuring that the captured chromosomes dynamics approaches the natural features of meiosis. Here, we report that *Arabidopsis thaliana* flowers cultivated in a culture medium (CCM) composed of the half-strength Murashige and Skoog basal salt, MES, Myo-inositol, sucrose and agar produce diploid microspores due to occurrence of meiotic restitution. Cytological studies revealed adjacent nuclei distribution and incomplete cytokinesis at late meiosis II in meiocytes within the CCM flowers. Immunolocalization of α-tubulin and the microtubule-associated protein MAP65-3 showed that the orientation of spindles at metaphase II and the organization of radial microtubule arrays at the tetrad stage are interfered, which explains the production of meiotically-restituted microspores. Moreover, the CCM flowers showed a gradually impaired expression of Aborted Microspores (AMS), a key transcription factor regulating tapetum development and meiotic cytokinesis. Interestingly, an increased supply of sucrose in culture medium promoted the expression of AMS and partially rescued haploid microspore formation in the CCM flowers. Taken together, this study suggests a role of sucrose in facilitating meiotic cytokinesis and gametophytic ploidy stability in plants.

**One-sentence summary:** Arabidopsis flowers cultivated in culture medium produce unreduced microspores due to interfered meiotic cytokinesis, which is partially rescued by increased sucrose supply.

## Introduction

Meiosis is a specialized-type of cell division in which chromosomal DNA is replicated once followed by twice nuclei division. Meiosis leads to genetic diversity through recombination of homologous chromosomes and production of gametes with a halved chromosome number, which are required for genome stability over generations (Zickler and Kleckner, 2023). Most angiosperms have experienced whole genome duplication (WGD) or polyploidization (WGP) in their evolution history, which plays an important role in speciation and environmental adaption (Otto, 2007; Ren et al., 2018; Van de Peer et al., 2020). Formation of unreduced gametes through meiotic restitution, a phenomenon that defines non-reductional meiosis events, is considered the main route to WGD in flowering plants (Ramsey and Schemske, 1998).

In flowering plants, defects occurring in multiple meiosis processes can lead to formation of unreduced gametes. In Arabidopsis, potato and horticultural plant species, dysfunction or down-regulation of the spindle regulators *JASON* or *Parallel Spindle 1* (*PS1*) causes a parallel and/or triad-like configuration of spindles during meiosis II, leading to a failure in the separation of nuclei and thus production of diploid gametes and polyploid offspring (Andreuzza et al., 2015; Clot et al., 2024; d’Erfurth et al., 2008; De Storme and Geelen, 2011; Peloquin et al., 1999; Zhou et al., 2022b). Omission of meiotic cell cycle triggered by functional defects in the meiotic cell cycle regulators *Omission of Second Division 1* (*OSD1*) and *Tardy Asynchronous Meiosis* (*TAM*/*CYCA1;2*) (d’Erfurth et al., 2010; d’Erfurth et al., 2009; Pang et al., 2025; Zhou et al., 2022b) induces meiotically-restituted dyads and thus diploid gametes in multiple species. Moreover, irregular cytokinesis interferes with chromosome distribution and thus can also induce unreduced gamete formation (Liu et al., 2021a; Spielman et al., 1997; Takahashi et al., 2010; Yang et al., 2003; Zeng et al., 2011). In addition to the genetic lesions, multiple abiotic stresses, including extreme temperatures and ultraviolet radiation, have been reported to induce meiotic restitution in plants (De Storme and Geelen, 2020; Fu et al., 2024; Mai et al., 2019; Zhou et al., 2022a).

Live-imaging microscopy technology has been increasingly used in meiosis study in plants because of its advantage in obtaining insights into dynamic cellular processes (Prusicki et al., 2021). In this system, the dynamic behaviors of meiotic chromosomes and proteins are monitored in alive meiocytes wrapped in the anthers of the inflorescences *ex vivo* cultivated in established culture medium (Prusicki et al., 2019). Based on the differences in tissues, experimental purposes and species, different types of culture medium are used for the live-imaging analysis of meiosis, in which the Murashige and Skoog (MS) salt has been used as the basal component with additives for suitable applications on different microscopic systems (Prusicki et al., 2021). In the currently established culture systems, meiosis features including duration of meiosis and recombination rate recorded via the live-imaging technology are comparable to the findings obtained from fixed tissues, even under stressful environmental conditions (De Jaeger-Braet et al., 2022; France et al., 2021; Ning et al., 2021; Prusicki et al., 2019; Yang et al., 2022). Additionally, such an *ex vivo* system provides a convenient and reliable strategy for analyzing features of meiosis exposing to chemicals for biochemical and genetic evidences (Yuan et al., 2025). However, the potential impact of cultivating inflorescences in basal culture medium without abundant additives, such as vitamins, on meiosis has not been determined, which is relevant for developing reliable culture systems for different experimental purposes and in different plant species.

In this study, we report that Arabidopsis (*Arabidopsis thaliana*) flowers cultivated in a basal culture medium produce unreduced microspores due to mis-organization of microtubule cytoskeleton during meiosis II and thus a consequent restituted meiotic cell division. Moreover, we showed that an increased sucrose supply partially rescues tapetum development and formation of haploid microspores. Taken together, this study suggests a positive role of sucrose in facilitating meiotic cytokinesis and the stability of gametophytic ploidy in Arabidopsis.

## Results

### Arabidopsis flowers cultivated in culture medium produces diploid microspores

To explore the effect of cultivating inflorescences in culture medium without abundant additives on meiosis in plants, we cut inflorescences from young flowering Arabidopsis (*Arabidopsis thaliana*) and then cultivated them in a culture medium composed of the half-strength Murashige and Skoog basal salt, MES (0.05% [w/v]), Myo-inositol (0.01% [w/v]), sucrose (1% [w/v]) and agar (0.8% [w/v]). The flowers from the inflorescences cut from intact plants directly used for experiments were used as the control samples. The Arabidopsis *quartet 1* (*qrt1*) mutant, in which released microspores maintain a tetrad configuration (Francis et al., 2006), was used for the analysis of meiosis products. In control flowers, only normal tetrads that consisted of four haploid microspores that indicated normal meiosis were observed (Fig. 1A, B and E). In flowers cultivated in the culture medium (CCM) for 40∼48 h, we found about 2.4% pollen mother cells (PMCs) at the unicellular microspore stage exhibiting configurations of a triad, which harbored one diploid nucleus and two haploid nuclei, or a dyad that contained two equally-sized microspores with each showing two nuclei (Fig. 1C-E). These figures suggested that PMCs in Arabidopsis CCM flowers produce unreduced microspores likely resulting from occurrence of meiotic restitution.

**Fig. 1.**
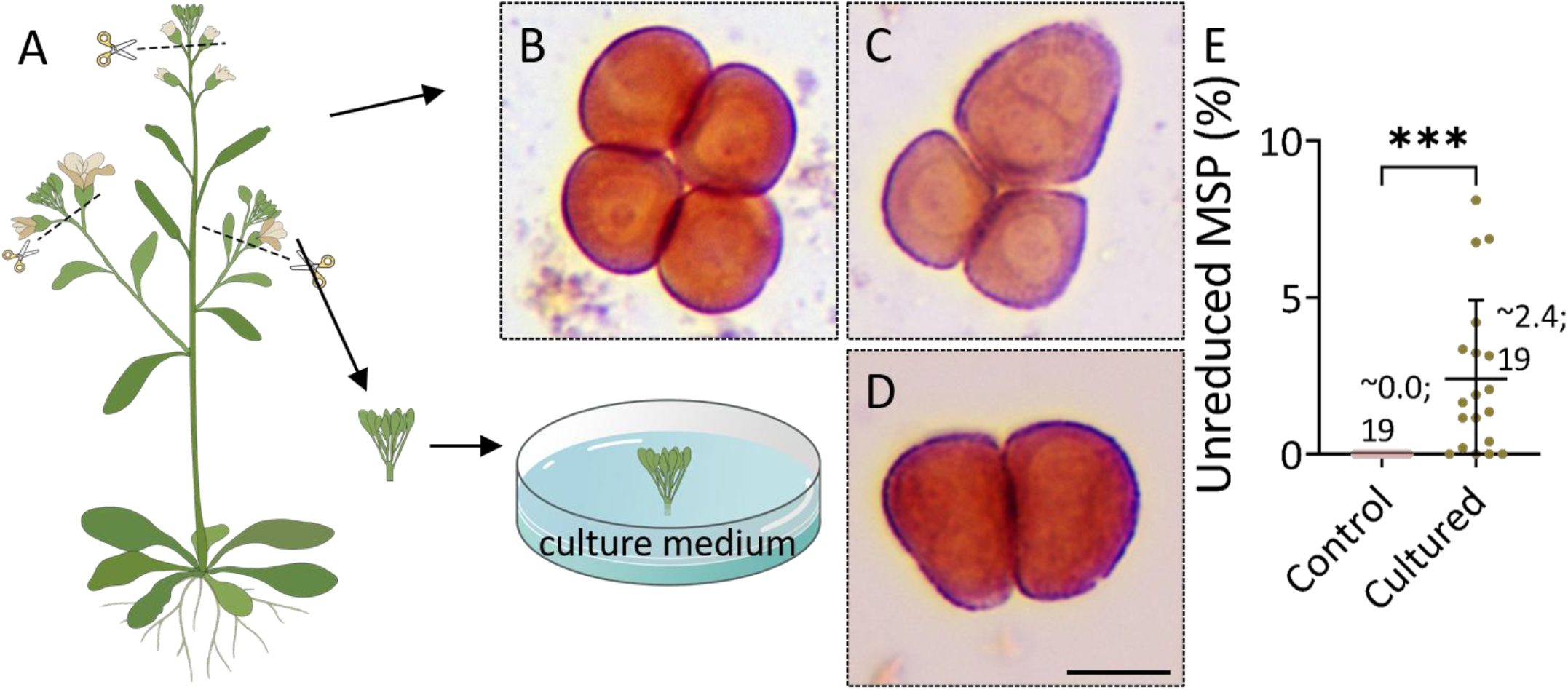
Arabidopsis flowers cultivated in culture medium produce diploid microspores. A, A model describing the manipulation of cultivating an Arabidopsis inflorescence in culture medium. B-D, A tetrad (B), a triad (C) and a balanced-dyad (D) at unicellular microspore stage produced by control or CCM flowers in the *qrt* mutant. E, Graph showing the rate of unreduced microspores yielded by control and CCM flowers. The significance level was determined based on an unpaired *t* test; the average rate of the unreduced microspores and the number of analyzed inflorescences are shown; *** indicates *P* < 0.001; CCM, cultivated in culture medium; MSP, microspore. Scale bar, 10 μm.

### PMCs in CCM flowers show defects in meiotic cytokinesis

To confirm the occurrence of meiotic restitution in PMCs in Arabidopsis CCM flowers, we stained PMCs at the tetrad and microspore stages in the *qrt* mutant with 4’,6-diamidino-2-phenylindole (DAPI). Tetrad-staged PMCs from control flowers showed separated haploid nuclei in corners of the cells, between which organelle bands were visible (Fig. 2A). These tetrads developed into unicellular microspores with each harboring a single haploid nucleus (Fig. 2C). In the CCM flowers, adjacent distribution of two nuclei was visualized at tetrad stage, and there was no organelle band between the adjacent nuclei, which showed a triad configuration (Fig. 2B). These triads resulted in production of microspores with two similarly-sized nuclei that indicated diploid microspores (Fig. 2D and E). At later developmental stages, triad microspores which showed stick cell walls that blocked entry of DAPI into cells with failed nuclei staining were observed (Fig. 2F).

**Figure 2.**
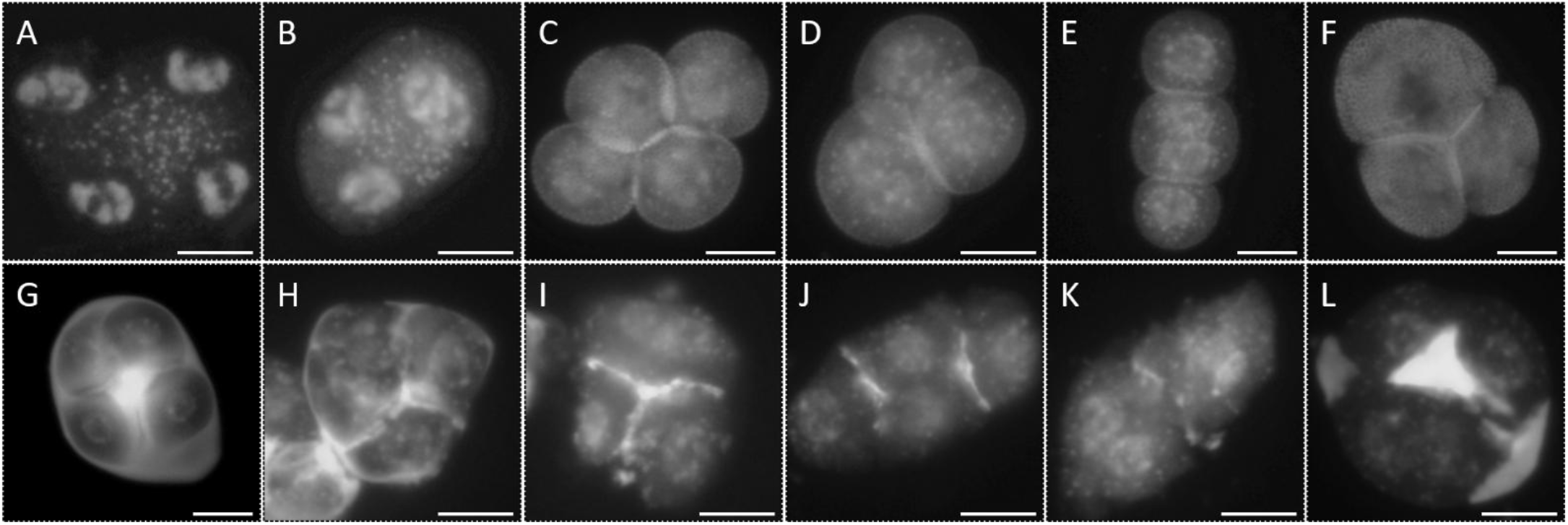
Pollen mother cells in flowers cultivated in culture medium show defects in meiotic cytokinesis. A and B, DAPI-staining of PMCs at the tetrad stage showing normal (A) and adjacent (B) nuclei distribution. C-F, DAPI-stained unicellular microspores observed in control (C) showing a normal tetrad configuration and flowers cultivated in culture medium (CCM) (D-F) showing a triad configuration. G-L, Combined DAPI and aniline blue staining of tetrad-staged PMCs in control (G) and CCM flowers (H-L). Scale bars, 10 μm.

The irregular distribution of haploid nuclei suggested that cell walls were not successfully built and thus indicated for a defect in cytokinesis. To this end, we stained tetrad-staged PMCs with DAPI and aniline blue, which labels callose, the main component of the meiotic cell walls during meiosis and at early microspore stages. In a tetrad, callosic cell walls were constructed between four isolated nuclei preventing them from fusing and thus facilitating formation of haploid microspores (Fig. 2G). Remarkably, tetrad-staged PMCs in CCM flowers exhibited failed and/or incomplete assembly of callosic cell walls, leading to an adjacent localization of nuclei (Fig. 2H-L). These observations revealed a defect in meiotic cytokinesis in PMCs from the CCM flowers.

### PMCs in CCM flowers show irregular chromosome distribution at the end of meiosis II

To further characterize the meiotic defects in Arabidopsis CCM flowers, we analyzed chromosome spreads by DAPI staining. In control flowers, meiocytes at pachytene stage displayed full juxtaposition of homologous chromosomes (homologs) (Fig. 3A) indicating complete homolog pairing and synapsis. Five bivalents were consistently observed at diakinesis and metaphase I (M I) (Fig. 3B and C), which manifested completion of meiotic recombination. Homologs were separated at anaphase I (A I) and temporally decondensed at interkinesis, which recondensed and were aligned at two cell polars at M II by the pulling force from two spindles (Fig. 3D-F). At telophase II (T II) stage, separation of sister chromatids leaded to formation of four haploid chromosome sets, with each developing into a haploid nucleus at the tetrad stage (Fig. 3G and H). In the CCM flowers, meiocytes during meiosis I did not show an obvious defect (Fig. 3I-M), which indicated regular meiotic recombination and homolog separation. However, defective orientation of aligned chromosomes at M II was visualized (Fig. 3N and O), which implied an improper organization and/or mis-orientation of spindles. At T II, an adjacent localization of nuclei indicative for irregular chromosome distribution was observed (Fig. 3P and Q), which leaded to the production of tetrad meiocytes showing a triad configuration (Fig. 3R-T). Hence, the observed meiotically-restituted unreduced microspores likely resulted from the defective distribution of chromosome sets and nuclei at the end of meiosis II.

**Fig. 3.**
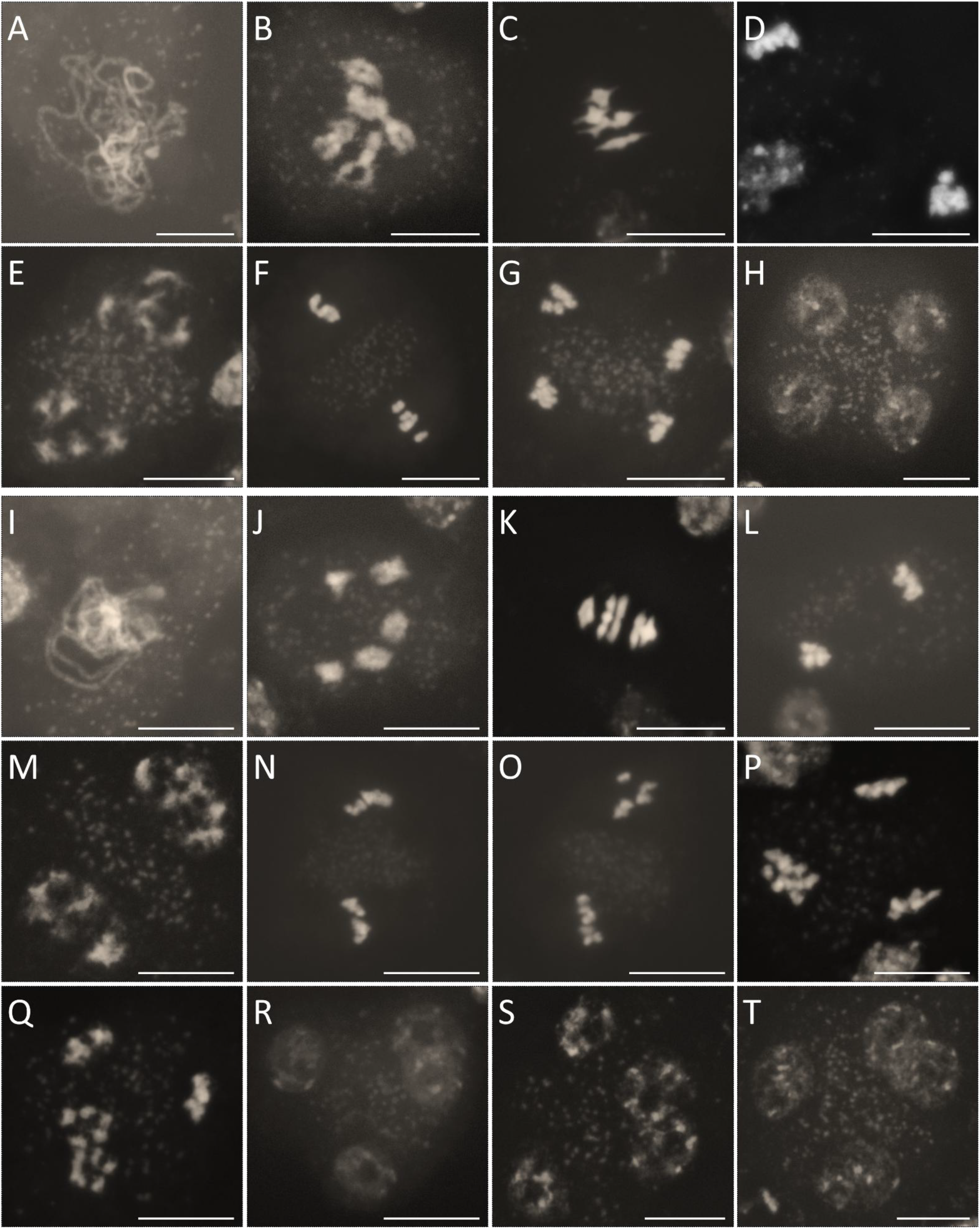
PMCs in flowers cultivated in culture medium show adjacent nuclei distribution at the end of meiosis II. A-T, DAPI-stained meiotic chromosome spreads at pachytene (A and I), diakinesis (B and J), metaphase I (C and K), anaphase I (D and L), interkinesis (E and M), metaphase II (F, N and O), telophase II (G, P and Q) and tetrad (H and R-T) stages in control flowers (A-H) and the flowers cultivated in culture medium (I-T). Scale bars, 10 μm.

### PMCs in CCM flowers show irregular spindle and phragmoplast organization at meiosis II

To determine whether the lesions in meiotic chromosome distribution and cytokinesis in CCM flowers are induced by defects in microtubular cytoskeleton, we analyzed organization of spindles and phragmoplasts by performing immunolocalization of tubulin and the microtubule-associated protein MAP65-3 using an anti-α-tubulin and an anti-GFP antibodies in the *pMAP65-3::MAP65-3-GFP* reporter line (Sofroni et al., 2020). In meiocytes from both control and CCM flowers, a spindle was built at M I with the MAP65-3 protein co-localizing with microtubules (Fig. 4A). At A I and interkinesis, a phragmoplast structure was organized between separated homologs, and MAP65-3 localized at the middle region of the phragmoplast (Fig. 4B and C). These data confirmed that the CCM flowers have no defect in meiosis I. In control meiocytes at M II, two spindles were perpendicularly organized to drive the segregation of sister chromatids at T II, and MAP65-3 displayed a similar localization pattern as that in meiosis I (Fig. 4D and E). At the tetrad stage, mini-phragmoplasts composed of radial microtubule arrays (RMAs) were assembled between isolated haploid nucleus, and MAP65-3 localized at the center of the mini-phragmoplasts (Fig. F). In the CCM flowers, the localization pattern of MAP65-3 was not altered (Fig. 4G-P). However, meiocytes at M II and T II stages, respectively, displayed a triangle configuration of spindles and phragmoplasts (Fig. 4H-I), indicating an irregular orientation. At the tetrad stage, triads showing an adjacent localization of nuclei and an omission of RMAs were visualized (Fig. 4J-N). Moreover, in some tetrads that showed separated haploid nuclei, RMAs and MAP65-3 were not regularly assembled between the nuclei (Fig. 4O and P). The defective microtubule assembly and/or organization could explain the lesions in meiotic chromosome distribution and cytokinesis in the CCM flowers.

**Fig. 4.**
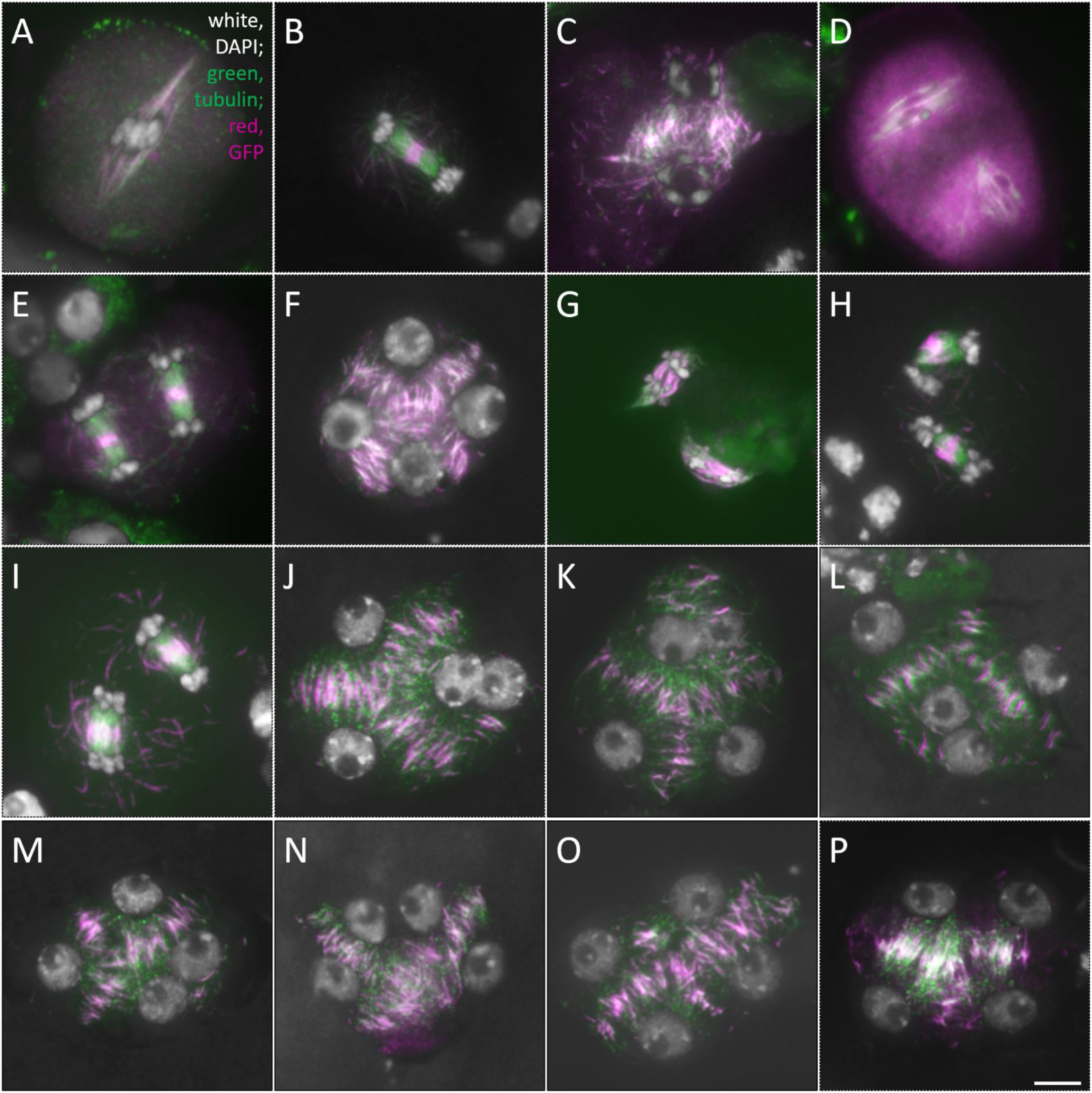
Meiocytes in flowers cultivated in culture medium show irregular microtubule organization during meiosis II. A-P, Immunolocalization of alpha-tubulin (green) and MAP65-3-GFP (red) in meiocytes at metaphase I (A), anaphase I (B), interkinesis (C), metaphase II (D and G), telophase II (E, H and I) and tetrad (F, J-P) stages showing normal (A-F) or abnormal organization and/or assembly (G-P) observed in control flowers and flowers cultivated in culture medium. Scale bar, 10 μm.

### Increased sucrose supply partially rescues AMS expression in the CCM flowers

It has previously been reported that normal development of tapetum is needed for faithful organization of RMAs and meiotic cytokinesis in Arabidopsis (Tidy et al., 2022). The lesions in meiotic cell wall formation and RMA organization in meiocytes led us to hypothesize that the meiotic restitution in CCM flowers is triggered by an attenuated function and/or development of the tapetum. To this end, we analyzed the expression of *Aborted Microspores* (*AMS*), a transcription factor required for tapetum development, dysfunction of which induces defective meiotic cytokinesis and thus meiotic restitution, by performing live-imaging using the CCM flowers from a *pAMS::AMS-GFP* reporter (Xiong et al., 2016). In control flowers, anthers at the tetrad and unicellular microspore stages showed expression of AMS-GFP specifically in the tapetal cell layer (Fig. 5A and B). In the tetrad-staged anthers at one-day post cultivation (1 dpc) in the medium, there was no obvious alteration detected in the expression of AMS-GFP (Fig. 5C). However, reduced AMS-GFP expression was visualized in the anthers at 2 dpc, and no AMS-GFP was detected at 3 dpc (Fig. 5E and G). These data indicated that the expression of AMS is damaged in the CCM flowers.

**Fig. 5.**
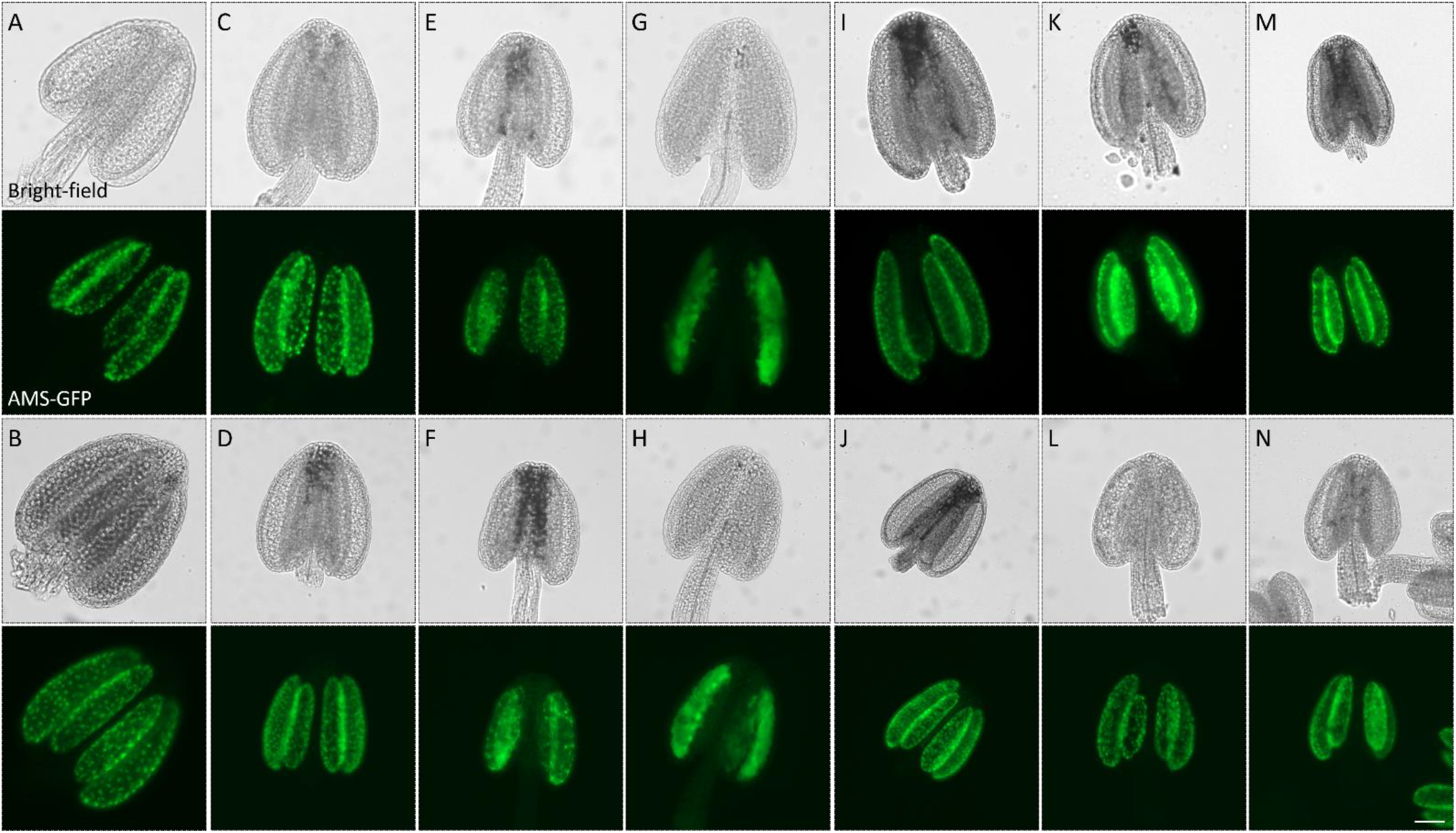
Increased sucrose or fructose supply rescues AMS expression in the tapetum. A and B, Expression of AMS-GFP in the tapetum of anthers at the tetrad (A) and microspore (B) stages in control flowers. C-H, Expression of AMS-GFP in the tapetum of anthers at the tetrad stage in flowers cultivated in culture medium containing 1% sucrose (C, E and G) or 1% fructose (D, F and H) sucrose for 24 (C and D), 48 (E and F) and 72 h (G and H). I-N, Expression of AMS-GFP in the tapetum of anthers at the tetrad stage in flowers cultivated in culture medium containing 10% sucrose (I, K and M) and 10% fructose (J, L and N) for 24 (I and J), 48 (K and L) and 72 h (M and N). Scale bar, 50 μm.

It has been reported that faithful tapetum development relies on normal sugar metabolism in the flowers (Borghi, 2025; Liu et al., 2021b). We wondered whether the damaged AMS expression in the CCM flowers is due to a shortage of sucrose supply in the anthers. Therefore, we monitored the expression of AMS in the CCM flowers exposed to an elevated sucrose concentration. In most flowers from 1 to 3 dpc, we observed normal expression of AMS-GFP in the anthers at the tetrad stage, which indicated that the increased sucrose supply partially rescued the expression of AMS in the tapetum (Fig. 5I, K and M). Because sucrose breaks down into other sugars to induce downstream cellular responses (Yoon et al., 2021), we tested whether fructose, a metabolite of sucrose, has a similar effect on the expression of AMS in the CCM flowers. The CCM flowers exposed to 1% fructose showed normal AMS expression at 1 dpc, but exhibited a reduced and an impaired AMS expression at 2 and 3 dpc, respectively (Fig. 5D, F and H). Under a 10% fructose condition, the expression of AMS in most flowers at 2 and 3 dpc was recovered (Fig. 5J, L and N). These findings reveal a positive impact of sucrose and its metabolite on AMS expression in the anthers.

### Increased sucrose supply partially rescues haploid microspore formation

To test whether an increased sucrose supply could complement the meiotic cytokinesis defects in the CCM flowers, we quantified the rates of unreduced microspores in the CCM flowers exposed to different concentrations of sucrose. In control flowers, only tetrads and haploid microspores were observed (Fig. 6A, E and I). In flowers cultivated in culture medium with 1% sucrose, meiocytes showing meiotic restitution at the tetrad stage was visualized and about 3.1% unreduced microspores were recorded (Fig. 6B-D, F-H and I). A similar rate of unreduced microspores was found in the flowers grown in medium without sucrose as that in 1% sucrose-supplied medium (Fig. 6I). Interestingly, flowers cultivated in medium with 10% sucrose yielded a significantly lower rate (∼1.3%) of unreduced microspores (Fig. 6I, *P* < 0.01), which suggested a partially rescued meiotic cytokinesis and haploid microspore formation.

**Fig. 6.**
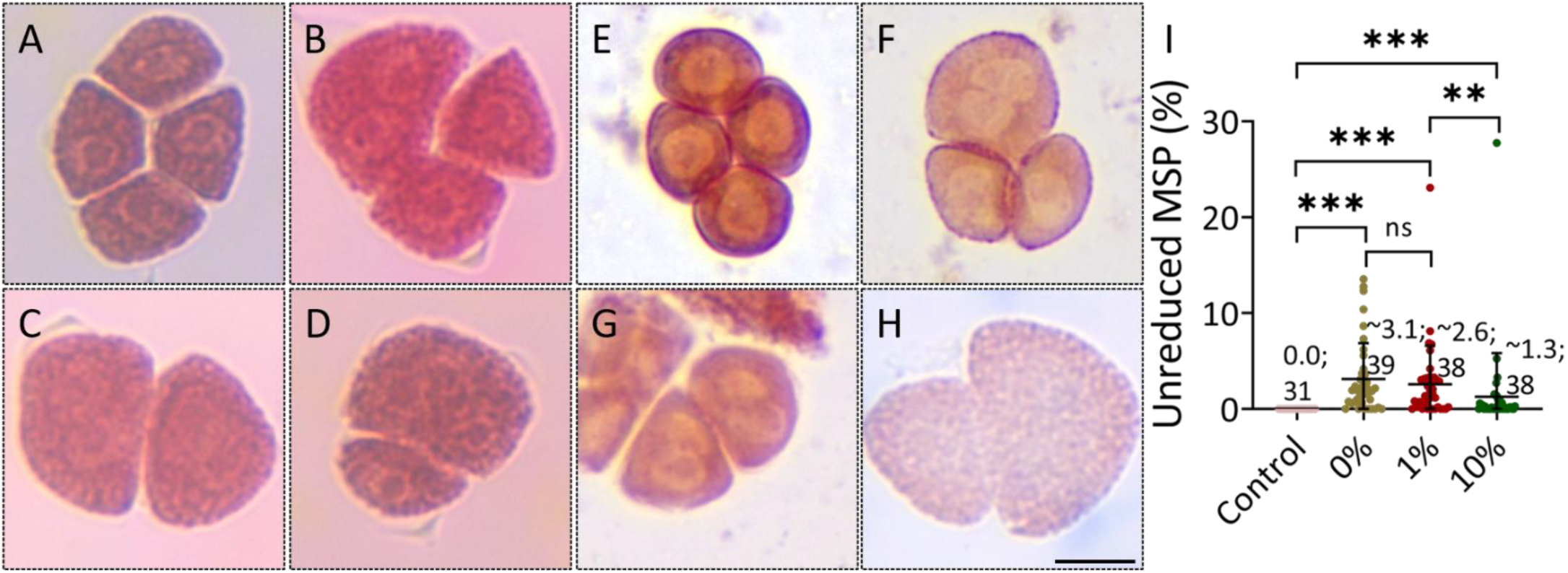
Increased sucrose supply in culture medium partially reduces diploid microspore formation. A-H, Orcein-staining of meiocytes at tetrad stage (A-D) and unicellular microspores (E-H) in control flowers (A and E) and flowers cultivated in culture medium (B-D, F-H) showing a tetrad (A and E), a triad (B and F), a balanced-dyad (C and G) or an unbalanced-dyad (H) configuration. I, Graph showing the rate of diploid microspores yielded by control flowers and flowers in culture medium containing 0%, 1% and 10% sucrose, respectively. Significance levels were determined based on unpaired *t* tests; the average rate of unreduced microspores and the number of inflorescences are shown; *** indicates *P* < 0.001; ** indicates *P* < 0.01; ns indicates *P* > 0.05; MSP, microspore. Scale bar, 10 μm.

## Discussion

In this study, we showed that Arabidopsis flowers cultivated in culture medium produce unreduced microspores due to occurrence of meiotic restitution caused by defective meiotic cytokinesis. Further cytological studies revealed that the orientation of spindles and phragmoplasts during meiosis II is altered, which results in irregular formation and/or organization of RMAs at the tetrad stage in the CCM PMCs. The altered orientation of spindles and phragmoplast at meiosis II in the CCM flowers is possibly owing to an interfered expression and/or function of the spindle regulator JASON and/or AtPS1 (De Storme and Geelen, 2011), which have been proposed to mediate the response of microtubule organization to environmental stimulus (Cabout et al., 2017; Fu et al., 2024). However, the tetrads that showed normal nuclei distribution but failed assembly of RMAs and MAP65-3 localization between the separated nuclei (Fig. 4P) suggest that the formation and/or composition of the organelle bands, which act as physical barriers between metaphase II spindles and thus ensure correct chromosome distribution (Brownfield et al., 2015), may be attenuated independently of the impacted JASON function (Gasser et al., 1988; Koç and De Storme, 2022).

The occurrence of meiotic restitution and the microtubule organization defects have not been reported in previous studies that cultured Arabidopsis inflorescences in growth medium for live-imaging microscopy (Prusicki et al., 2019; Sofroni et al., 2020; Valuchova et al., 2022; Wijnker et al., 2019; Yang et al., 2020; Yang et al., 2022; Yuan et al., 2025). The culture medium used in this study only contained Myo-inositol yet without other vitamins, such as nicotinic acid, pyridoxine hydrochloride, glycine and thiamine hydrochloride, which play important roles in metabolic regulation including energy metabolism, amino acid synthesis, oxidation-reduction reactions, stress response and cell development in plants (Berglund et al., 2017; Schnellbaecher et al., 2019; Sultana et al., 2019). Thus, we speculate that the meiosis defects we observed are induced by cellular lesions due to a lack of the chemicals necessary for the development of meiocytes (Prusicki et al., 2021). Hence, the components and possibly also their dosages in the culture medium should be tested and delicately modified for the establishment of an *ex vivo* culture system for plant meiosis study. On the other hand, the average frequency of the unreduced microspores in the CCM flowers is quite low (less than 3%), which plausibly makes it difficult to capture the lesions under the fast-moving and dynamic intracellular conditions. Considering the complexity of the chromosome dynamics during meiosis especially during recombination, minor differences may exist between the features of meiosis captured in flowers grown on the *ex vivo* culture system with those under natural conditions.

It has been reported that the normal development of the tapetum at early flower stages is needed for successful RMA organization and the assembly and regular configuration of cytokinetic cell walls during meiosis in Arabidopsis (Liu et al., 2017; Tidy et al., 2022). The defects in callosic cell walls together with the impairment of AMS expression in the anthers suggest that the meiotic restitution in the CCM flowers could be caused by damaged development and function of the tapetum, which thereafter attenuates the expression of meiosis genes or the metabolism of enzymes and components for organelle band formation and cell wall assembly in meiocytes (Biswas and Chaudhuri, 2024; Lei and Liu, 2020; Muro et al., 2025; Wei and Ma, 2023). Whether and how the tapetum plays a role in regulating the expression and/or function of JASON and AtPS1, or other microtubule regulators, await further studies. We found that an increased concentration of sucrose in the culture medium promoted AMS expression and haploid microspore formation, which could be attributed to a compensated energy supply or activation of signaling pathways mediated by sucrose and/or its metabolites in the anthers (Borghi, 2025; Liu et al., 2021b; Wang et al., 2022; Yoon et al., 2021). It can be noted that the rate of unreduced microspores largely varies between individual CCM inflorescences (Fig. 6I), which may result from the difference in the inherent sucrose abundance in those inflorescences. In multiple plant species, the regulatory modules of the tapetum development and meiosis progression are tightly associated with the factors in sugar metabolism and/or signalings (Lei and Liu, 2020; Liu et al., 2021b; Sun et al., 2025; Wang et al., 2023). In addition, in rice, the TDR Interacting Protein 2 (TIP2), a basic helix-loop-helix protein required for tapetum development, regulates the expression of carbohydrate-active glycosyltransferases and glycosyl hydrolases in the tapetum, dysfunction of which has been found to cause arrested meiosis progression (Fu et al., 2014; Wang et al., 2025). These findings suggest that the tapetum may influence the fidelity of meiotic cytokinesis by regulating sugar metabolism. We propose that the partially rescued microsporogenesis by increased sucrose supply is a consequence of the recovered tapetum function.

Overall, our study reveals a role of sucrose in facilitating tapetum development and meiotic cytokinesis. Moreover, cultivating flowers in medium with modifications of components including sugars for ensuring anther development could be potentially developed as a strategy for inducing unreduced gametes in polyploid breeding programs.

## Material and methods

### Plant materials and growth conditions

*Arabidopsis thaliana* mutant *quartet* (*qrt*) (Francis et al., 2006), the *pMAP65-3::GFP-MAP65-3* (Sofroni et al., 2020) and *pAMS::AMS-GFP* (Xiong et al., 2016) reporters were used in this study. Seeds were germinated in soil for 6-8 days and seedlings were transferred to soil and cultivated in growth chambers with a 16 h day/8 h night, 20°C, and 50% humidity condition. For flower cultivation, young inflorescences were cut with the tips being embedded into the medium. A suitable amount of distilled water was added to the medium to avoid drying of the inflorescences.

### Preparation of culture medium

The culture medium used in this study was prepared by dissolving the basal MS salt (0.5 X), MES (0.05% [w/v]), Myo-inositol (0.01% [w/v]), sucrose (1% [w/v]) and agar (0.8% [w/v]) in distilled water with a pH at 5.7.

### Cytological analysis of meiocytes and microspores

Orcein staining, 4,6-diamidino-2-phenylindole (DAPI) staining, and aniline blue staining of meiocytes were performed as described previously (Fu et al., 2024). Quantification of microspores was performed at 48 h post the cultivation of flowers in the medium.

### Preparation of chromosome spreads

Inflorescences of young Arabidopsis were fixed in precooled Carnoy’s fixative for at least 24 h. Meiosis-staged flower buds were washed twice with distilled water and once with citrate buffer (10 mM, pH = 4.5), followed by incubation in a digestion enzyme mixture (0.3% pectolyase and 0.3% cellulase in citrate buffer, 10 mM, pH = 4.5) at 37°C for 2.5h. Subsequently, the digested flower buds were washed once in distilled water and macerated in distilled water on a glass slide. Two aliquots of 60% acetic acid were added to the slide, which was dried on a hotplate at 45°C for 2 min. The slide was washed with ice-cold Carnoy’s fixative and then air-dried. DAPI (5 μg/mL) diluted in antifade mounting medium was added to the slide, and the coverslip was mounted and sealed with nail polish.

### Immunolocalization of microtubules and MAP65-3 protein

Immunolocalization assays were performed by referring to (Liu et al., 2017; Wang et al., 2014). The antibodies against a-tubulin (Lei et al., 2020) and GFP (Zhao et al., 2023) were diluted by 1:500 and 1:300, respectively. The secondary antibodies have been described previously (Lei et al., 2020; Zhao et al., 2023).

### Live-imaging of reporters

To analyze the expression of AMS, anthers in flowering *pAMS::AMS-GFP* reporter were isolated and placed on a glass slide with a drop of distilled water being added to the samples, which was then mounted with a cover slide and examined under an inverted fluorescence microscope. The developmental stages of the anthers were determined by checking the released PMCs or microspores in the anthers.

### Microscopy

Fluorescence microscopy was performed using an Olympus IX83 inverted fluorescence microscope equipped with an X-Cite lamp and a Prime BSI camera. Bifluorescent images and Z-stacks were processed using Image J.

### Statistical analysis

Significance analysis was performed using the unpaired *t*-test with GRAPHPAD PRISM (v.8), and significance level was set as *P* < 0.05. Variation bars indicate SD.

## Funding

This study was supported by Hubei Provincial Natural Science Foundation of China (2024AFB695), the Fundamental Research Funds for the Central Universities, South-Central Minzu University (CZZ24011), Zhejiang Science and Technology Major Program on Agricultural New Variety Breeding (2021C02071–6 to Z.R.), Zhejiang Provincial Natural Science Foundation (ZCLTGN24C1601 to Z.R.) and Zhejiang Sci-Tech University Start-up Fund (22052138-Y to Z.R.).

## Data availability statement

The data that support the findings of this study are available from the corresponding author (B.L.) upon reasonable request.

## CRediT authorship contribution statement

H.F., Y.C., X.C., H.H. and J.W. contributed to investigation; C.W. and Z.R. contributed to data analysis; B.L. conceived project, analyzed data, wrote, and edited the manuscript. All the authors have read and agreed with the manuscript prior to the submission.

## Declaration of Competing Interest

The authors declare that they have no known competing financial interests or personal relationships that could have appeared to influence the work reported in this paper. The authors declare the following financial interests/personal relationships which may be considered as potential competing interests.

## Acknowledgement

The authors thank Arp Schnittger (Universität Hamburg) for sharing the *pMAP65-3::MAP65-3-GFP* reporter; they also thank Zhongnan Yang (SJTU) and Yue Lou (SHNU) for sharing the *pAMS::AMS-GFP* reporter.

## Notes

### Competing Interest Statement

The authors have declared no competing interest.

